# Exploiting evolutionary non-commutativity to prevent the emergence of bacterial antibiotic resistance

**DOI:** 10.1101/007542

**Authors:** Daniel Nichol, Peter Jeavons, Alexander G. Fletcher, Robert A. Bonomo, Philip K. Maini, Jerome L. Paul, Robert A. Gatenby, Alexander R.A. Anderson, Jacob G. Scott

**Affiliations:** Department of Computer Science, University of Oxford, Oxford, UK; Wolfson Centre for Mathematical Biology, Mathematical Institute, University of Oxford, Oxford, UK; Department of Medicine, Louis Stokes Department of Veterans Affairs Hospital, Cleveland, OH, USA; School of Computing Sciences and Informatics, University of Cincinnati, Cincinnati, OH, USA; Department of Integrated Mathematical Oncology, H. Lee Moffitt Cancer Center and Research Institute, Tampa, FL, USA

**Keywords:** theoretical biology, antibiotic resistance, bacteria, mathematical model, evolution, fitness landscape, strong selection weak mutation

## Abstract

The increasing rate of antibiotic resistance and slowing discovery of novel antibiotic treatments presents a growing threat to public health. Here, we develop a Markov Chain model of evolution in asexually reproducing populations which is an encoding of the Strong Selection Weak Mutation model of evolution on fitness landscapes. This model associates the global properties of the fitness landscape with the algebraic properties of the Markov Chain transition matrix and allows us to derive general results on the non-commutativity and irreversibility of natural selection as well as antibiotic cycling strategies. Utilizing this formalism, we analyse 15 empirical fitness landscapes of E. coli under selection by di.erent beta-lactam antibiotics and demonstrate that the emergence of resistance to a given antibiotic can be both hindered and promoted by di.erent sequences of drug application. Further, we derive optimal drug application sequences with which we can probabilistically ‘steer’ the population through genotype space to avoid the emergence of resistance. This suggests a new strategy in the war against antibiotic.therapy.resistant organisms: drug sequencing to shepherd evolution through genotype space to states from which resistance cannot emerge and by which to maximize the chance of successful therapy.

**Background:** The increasing rate of antibiotic resistance and slowing discovery of novel antibiotic treatments presents a growing threat to public health. Previous studies of bacterial evolutionary dynamics have shown that populations evolving on fitness landscapes follow predictable paths. In this article, we develop a general mathematical model of evolution and hypothesise that it can be used to understand, and avoid, the emergence of antibiotic resistance.

**Methods and Findings:** We develop a Markov Chain model of evolution in asexually reproducing populations which is an encoding of the Strong Selection Weak Mutation model of evolution on fitness landscapes. This model associates the global properties of the fitness landscape with the algebraic properties of the Markov Chain transition matrix and allows us to derive general results on the non-commutativity and irreversibility of natural selection as well as antibiotic cycling strategies. Utilizing this formalism, we analyse 15 empirical fitness landscapes of *E. coli* under selection by different β-lactam antibiotics and demonstrate that the emergence of resistance to a given antibiotic can be both hindered and promoted by different sequences of drug application. We show that resistance to a given antibiotic is promoted in 61.4%, 68.6% and 70.3% of possible orderings of single, pair or triple prior drug administrations, respectively. Further, we derive optimal drug application sequences with which we can probabilistically ‘steer’ the population through genotype space to avoid the emergence of resistance.

**Conclusions:** Our model provides generalisable results of interest to theorists studying evolution as well as providing specific, testable predictions for experimentalists to validate our methods. Further, these results suggest a new strategy in the war against antibiotic-therapy-resistant organisms: drug sequencing to shepherd an evolving population through genotype space to states from which resistance cannot emerge and from which we can maximize the likelihood of successful therapy using existing drugs. While our results are derived using a specific class of antibiotics, the method is generalisable to other situations, including the emergence of resistance to targeted therapy in cancer and how species could change secondary to changing climate or geographical movement.

## Introduction

Resistance to antibiotic treatments within bacterial pathogens poses an increasing threat to public health, which coupled with the slowing discovery of novel antibiotics, could soon reach crisis point [Spellberg et al., 2013, French, 2010]. Novel classes of antibiotics discovered since 1987 are few in number [Silver, 2011] . Thus, it is becoming ever clearer that if we are to combat highly resistant bacterial infections, then we must find new ways to prevent resistance and new applications of existing antibiotics to these pathogens. Indeed, public health efforts have attempted to stem the emergence of resistance by reducing unnecessary prescription of antibiotics [Shlaes et al., 1997, Bartlett, 2011, Leuthner and Doern, 2013] and stopping the addition of sub-therapeutic antibiotics in livestock feed [Mathew et al., 2007]. However, such policies require global adoption to be truly effective [Moody et al., 2012], which they have not yet achieved, and is likely infeasible. Recently, there have been efforts to explore how existing antibiotics can be used in new ways to provide effective treatments for resistant pathogens, for example through combination therapy [Chait et al., 2007] or through cycling strategies [Goulart et al., 2013, Pena-Miller and Beardmore, 2010, Bergstrom et al., 2004] – long term hospital-scale treatment protocols that cycle which antibiotics are prescribed over timescales of weeks, months or years. However, our understanding of the mechanisms underlying these strategies remains limited. Indeed, from a mathematical model Pena-Miller and Beardmore [2010] predict that cycling strategies have the potential to perform both better and worse than mixed strategies and hence tools to find optimal strategies are needed.

In order to understand how to minimize the emergence of resistant pathogens, and to decide how best to treat them, we must first understand how their evolution is driven by the selective pressures of different antibiotic drugs — a fundamental problem of biology [de Visser and Krug, 2014]. In particular, if we understand which traits are likely to be selected for by which treatments, then we may be able to avoid selecting for those traits which confer resistance. Recent insights into the evolutionary process have yielded some actionable information. Specifically, Weinreich et al. 2005, Weinreich et al. 2006] showed that if the genome of a pathogen exhibits sign epistasis, where a given mutation is beneficial on some genetic backgrounds and deleterious on others, then there can exist inaccessible evolutionary trajectories. Further, Tan et al. [2011] studied the evolutionary trajectories of *Escherichia coli* under different antibiotics and found that adaptive mutations gained under one antibiotic are often irreversible when a second is applied. These irreversible trajectories can occur when resistance conferring mutations for one environment carry a cost in another which can be mitigated by other compensatory mutations [zur Wiesch et al., 2010, Tanaka and Valckenborgh, 2011]. These findings lead us to hypothesize that one antibiotic could be used to irreversibly steer the evolution of a population of pathogens to a genotype (or combination of genotypes) which is sensitive to a second antibiotic but also from which it is unlikely to acquire resistance to that antibiotic. This hypothesis was partly verified by the work of Imamovic and Sommer [2013] who demonstrated that evolving *E. coli* to become resistant to certain antibiotics can increase sensitivity to others. However, this work is limited in two ways. Firstly, in that the experiments do not exhaustively consider all evolutionary trajectories but instead highlight only those that arose in a small number of replicates and secondly, they do not consider how evolution will proceed once the second drug is applied and whether resistance can then emerge.

In this paper we present a Markov Chain model of evolution which we use to derive general results on the non-commutavity of evolution and requirements for cycling strategies. Then, using previously measured landscapes for 15 P-lactam anti-biotics, we illustrate that selective pressures are non-commutative, and that the emergence of resistance can be both hindered and promoted by different orderings of these pressures. These findings suggest new treatment strategies which use rational orderings of collections of drugs to shepherd evolution through genotype space to a configuration which is sensitive to treatment, as in the work of Imamovic and Sommer [2013], but also from which resistance cannot emerge.

## Evolution on Fitness Landscapes

We begin with the concept of a fitness landscape introduced by Wright [1932] and used by Weinreich et al. [2005] and Tan et al. [2011] to study evolutionary trajectories in asexually reproducing populations. We represent the genotype of an organism by a bit string of length N and model mutation as the process of flipping a single bit within this string. This model of mutation only accounts for point mutations and ignores the possibility of other biologically relevant mutations such as gene insertions, gene deletions and large structural changes to the genotype. This gives a set of 2^N^ possible genotypes in which individuals of a given genotype, say *g*, can give rise to mutated offspring which take genotypes given by one of the N mutational neighbors of *g* — precisely those genotypes g′ for which the Hamming distance [Hamming, 1950], Ham(*g,g′*), from *g* is 1. As such, our genotype space can be represented by an undirected N-dimensional (hyper-)cube graph with vertices in {0,1}^N^ representing genotypes and edges connecting mutational neighbors (Figure 1a).

We define a selective pressure on our graph that drives evolution, for example through an environmental change or drug application, as a fitness function

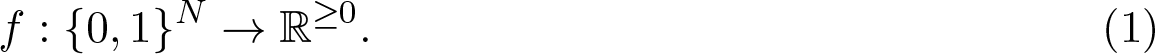

This fitness function represents a genotype-phenotype map in the simplest sense — assigning to each genotype a single real-valued fitness. Gillespie 1983, Gillespie 1984] showed that if the mutation rate u and population size *M* of a population satisfy *Mu* >> 1 >> *Mu*^2^, and if we assume that each mutation is either beneficial or deleterious, then each beneficial mutation in the population will either reach fixation or become extinct before a new mutation occurs. Further, selection will be sufficiently strong that any deleterious mutation will become extinct with high probability and hence we may assume that this always occurs. In the case that *Mu*^2^ ≈ 1 stochastic tunneling [Iwasa et al., 2004] through double mutations can occur and we cannot ignore deleterious mutations. Assuming Mu >> 1 >> Mu^2^, then after each mutation the population will stabilize to consist entirely of individuals with the same genotype and this genotype will be eventually replaced by a fitter neighboring genotype whenever one exists. This observation gives rise to the Strong Selection Weak Mutation (SSWM) model, which models a population as occupying a single vertex on a directed graph on the set of *2^N^* possible genotypes, {0,1}^N^, in which there exists an edge from vertex *a* to a neighboring vertex *b* if, and only if, *f (b)* > *f (a)* (see (Figure 1b and Figure 1c). This population undergoes a stochastic walk in which the population moves to an adjacent fitter genotype with some probability. Several ‘move rules’ have been proposed which can be used to select an adjacent fitter neighbor during this stochastic walk [Orr, 2005] and which of these move rules is most accurate depends on the population size [de Visser and Krug, 2014]. Common move rules include selecting the fittest neighbor [Kauffman and Levin, 1987, Fontana et al., 1993], selecting amongst fitter neighbors at random [Macken and Perelson, 1989, Macken et al., 1991, Flyvbjerg and Lautrup, 1992] or selecting fitter neighbors with probability proportional to the fitness increase conferred [Gillespie, 1983, Gillespie 1984, Gillespie 1991]. We encapsulate each of these variants of the SSWM model within our model.

## A Markov Model of Evolution

The SSWM model of evolution reduces the evolutionary process to a random walk on a directed graph and hence can be modeled by a Markov Chain [Grinstead and Snell, 1998]. For a fitness function 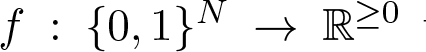 we can define a transition matrix 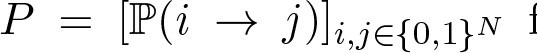 for a time-homogeneous absorbing Markov Chain by setting, for i ≠ j,

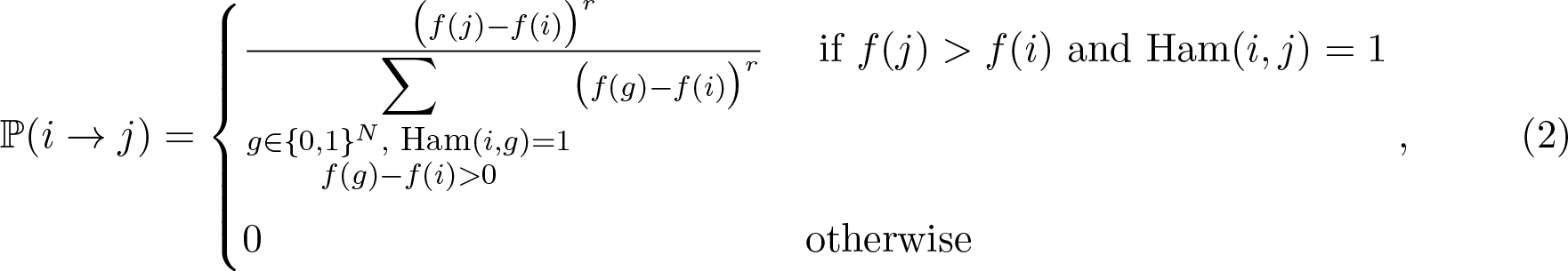

**Fig. 1.**
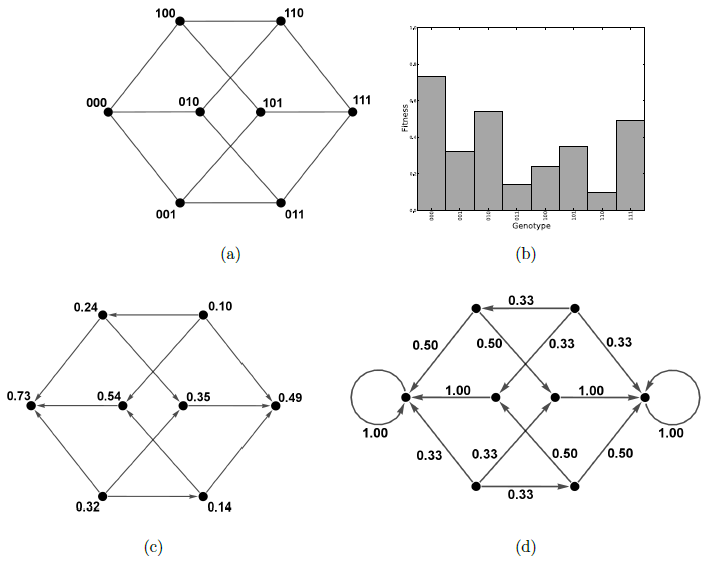
(a) The space of genotypes comprising bit strings of length *N* = 3. The vertices represent genotypes, and edges connect those genotypes which are mutational neighbors. (b) An example fitness landscape. (c) The directed evolutionary graph according to the landscape in (b) where the vertices still represent genotypes but are labeled by the associated fitness for clarity. The directed graph edges are determined by the fitness function and represent those mutations which can fix in a population (those which confer a fitness increase). (d) The Markov Chain constructed for the same landscape according to (Equation 2) and (Equation 3) with *r* = 0.

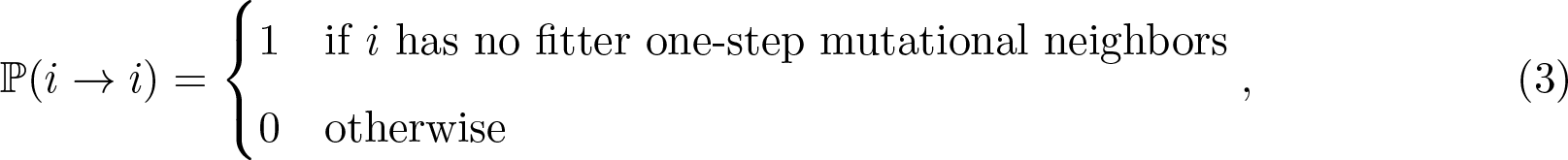

for each i (see Figure 1d. Here the parameter *r* ≥ 0 determines the extent to which the fitness increase of a mutation affects its likelihood of determining the next population genotype. In the case *r* = 0, we have the random move SSWM model (as in Macken and Perelson [1989], Macken et al. [1991], Flyvbjerg and Lautrup [1992]), in the limit r→ ∞ we have the steepest gradient ascent SSWM model (as in Kauffman and Levin [1987], Fontana et al. [1993]), and for *r* = 1 we have probability proportional to fitness increase (as in Gillespie 1983, Gillespie 1984, Gillespie 1991). This model differs from the Markov model used by Sella and Hirsh [2005] to study the neutral theory of evolution as we do not allow deleterious mutations to fix in the population.

Using this Markov Chain we can explore the possible evolutionary trajectories of a population on a given fitness landscape *f*. We next define a collection of population row vectors *μ^(t)^* for each *t ϵ ℕ*, where *μ^(t)^* has length *2^N^* and *k*^th^ component which gives the probability that the population has the *k*^th^ genotype at time *t* (where the genotypes are ordered numerically according to their binary value). These time steps *t* are an abstraction which discretely measure events of beneficial mutations occurring and fixing in the population. As such, the actual time between steps *t* and *t*+1 is not constant but may be considered drawn from a distribution parameterized by the mutation rate, reproductive rate and the number of beneficial mutations that can occur. This distribution could, for example, be determined by a Moran [Moran et al., 1962] or Wright-Fisher [Wright, 1932, Fisher, 1958—] process depending on how we choose to interpret the fitness values given by *f*. If the population has a genotype corresponding to a local optimum of the fitness landscape at time *t* then there are no beneficial mutations that can occur and this definition of a time step is not well defined. In this case there can be no more changes to the population under the SSWM assumptions and for mathematical convenience we define the probability of a local optimum population genotype remaining unchanged as one in equation 3 to ensure our model is a Markov Chain. In this case the step *t* to *t* + 1 can be chosen to take some fixed arbitrary time.

The distribution of a population at time *t* is related to its initial distribution, *μ*^(0)^, by

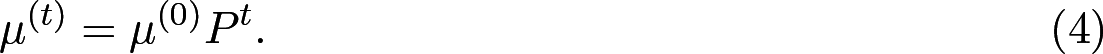

Since the Markov Chain is absorbing we know that there exists some *k* such that *P^k^P* = *P^k^* [Grinstead and Snell, 1998]. Consequently, we know that the matrix

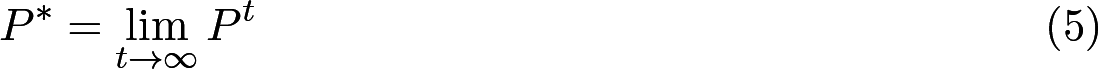

exists and in fact this limit is reached after only finitely many matrix multiplications. Thus a given initial population distribution μ^(0)^ will converge to a stationary distribution *μ^*^* after a finite number of steps in our model. Furthermore, if μ^*^ is known then we compute the stationary distribution μ^*^ as

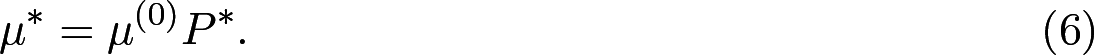

In particular, provided we assume a drug is applied for sufficiently long to ensure that the disease population reaches evolutionary equilibrium, we can explore the effects of applying multiple drugs sequentially by considering the matrices *P*^*^ for the associated fitness landscapes. In the following discussion we make this assumption.

By encoding the evolutionary dynamics in a Markov Chain we can investigate the evolutionary process from an algebraic perspective. In particular, as the transition matrix *P* encodes all of the evolutionary dynamics of the associated fitness landscape *f*, we can explore global properties of *f* by considering the algebraic properties of *P*. In the following section we present two simple, yet powerful, consequences of this observation.

## Non-Commutativity and Cycling of Natural Selection

We use the Markov Chain model to formally prove that for a large class of fitness landscape pairs, there is non-commutativity in the evolutionary process as described by the SSWM assumptions. More precisely, consider two drugs, *X* and *Y*, with corresponding fitness landscapes *x* and *y*. We wish to determine what, if any, difference there is between applying *X* followed by *Y* to a population as opposed to applying *Y* followed by *X* to that population. If we construct the transition matrices *P_x_* and *P_y_* corresponding to x and y, respectively, and take the limits 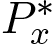 and 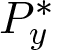 then our model predicts that the ordering makes no difference to the final population distribution on an initial population taking genotype *i* if, and only if,

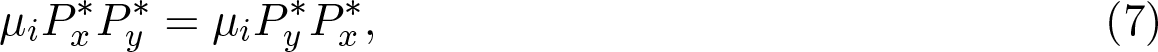

where μ_*i*_ is row vector of length 2^N^ whose *i*^th^ component is one and all of whose other components are zero.

If we do not know the starting population genotype we can only guarantee that the order of application is irrelevant when the outcome is the same regardless of the starting genotype. We require for all possible length 2^*N*^ unit vectors *μ_i_* that 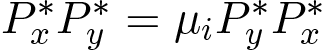. Since these unit vectors form a basis of ℝ^*N*^ this occurs precisely when

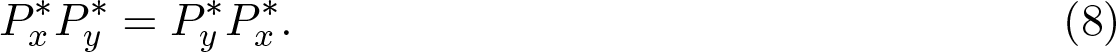

Hence drug application will only commute when the corresponding limit matrices commute. In practice we may be able to narrow down which genotypes are likely to constitute the population through bacterial genotyping or by observing that certain strains are not viable in the wild due to the high fitness cost of certain mutations. To determine how common commutativity is we first tested each pair of fitness landscapes from the landscapes empirically determined by Mira et al. [2014] for *E. coli* in the presence of N = 4 possible resistance conferring mutations under 15 different antibiotics (listed in Table 1). Of these 15 antibiotics we found no commuting pairs. We then tested 10^6^ pairs of random fitness landscapes with varying ruggedness generated according to Kauffman’s NK model for generating “tunably rugged” fitness landscapes [Kauffman and Levin, 1987, Kauffman and Weinberger, 1989] using a random neighborhood Boolean function for determining the fitness contributions of each locus. We fixed N = 5 and generated each landscape by first drawing *K* uniformly from {0,…,*N* − 1} and then using Kauffman’s model to build a landscape. We found that 0.132% of the landscape pairs generated had limit matrices which commuted, suggesting that commutativity is rare.

We now turn out attention to finding antibiotic cycling strategies as in Goulart et al. [2013]. Unless *x* is a flat landscape (taking equal values for all genotypes) there must exist a (not necessarily unique) genotype *j* whose fitness is a minimum and which has a fitter neighbor. Such a genotype satisfies ℙ[(*i* → *j*)] = 0 for all genotypes *i*. Hence if *x* is not flat, the limit matrix 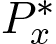 has at least one column of all zeros and is singular, so there cannot exist a second landscape *y* for which 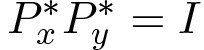. Hence there exists a unit row vector *μ_i_* for which 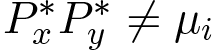 As the μ vectors encode probability distributions this means that natural selection in our model is irreversible in the sense that for a given (non-flat) landscape we cannot find another which is guaranteed to reverse its effects. This result precludes the existence of a general cycling strategy through which we can utilize a sequence of drugs to drive evolution through a cycle in genotype space such that the disease population returns to its original genotype, regardless of that starting genotype. If we do in fact know the starting genotype, as we might if the disease is contracted in the wild where resistance conferring mutations often carry a cost [Andersson and Hughes, 2010] making the wild-type genotype most likely, then cycling strategies can be found efficiently by our model. In particular, for a given starting genotype *i* the initial population distribution will be *μ_i_* and a sequence of drugs X_1_,⋯X_k_ with fitness landscapes *x_1_,⋯x_k_* will constitute a cycling strategy precisely when 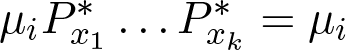. This criterion will be satisfied when μ_i_ is a left 1-eigenvector of 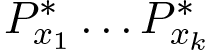. As such, we can find cycling strategies using matrix algebra and avoid the graph-search technique used by Goulart et al. [2013].

## Evolutionary Steering can both Prevent and Promote Resistance

Prescriptions of sequences of drugs occur frequently in the clinic, and often without any guidelines as to which orderings are preferable. Common examples of this include, but are not limited to, treatment of *H. pylori* [Gisbert et al., 2010], Hepatitis B [Hanazaki, 2004] and the ubiquitous change from broad to narrow spectrum antibiotics [Heenen et al., 2012]. The ordering of the sequence is therefore often determined arbitrarily, by the individual clinician's personal, or historical experience or from laboratory data. Ideally, we would like to be able to identify drug orderings that lower the probability of a highly resistant disease population emerging during the treatment. To consider optimal drug orderings in the context of our model we first need to know the fitness landscapes (or proxies of the fitness landscapes) of a number of antibiotics used to treat a given bacterial infection. Experimentally determining these landscapes requires one to consider all possible *2^N^* combinations of genotypes in a set of *N* genes, a task which is prohibitively difficult for all but small values of *N*. De Visser and Krug [2014] found that there have been less than 20 systematic empirical studies of fitness landscapes and that landscapes have been calculated for a number of model organisms including *E. coli* [Khan et al., 2011, Tan et al., 2011, Schenk et al., 2013, Goulart et al., 2013], *Saccharomyces cerevisiae* [Hall et al., 2010], *Plasmodium falciparum* [Lozovsky et al., 2009] and type 1 Human Immunodeficiency Virus [da Silva et al., 2010]. Recent work by Hinkley et al. [2011] which utilizes regression methods to approximate large fitness landscapes from samples of the space could help ameliorate the complexity of experimentally determining fitness landscapes.

**Fig. 2.**
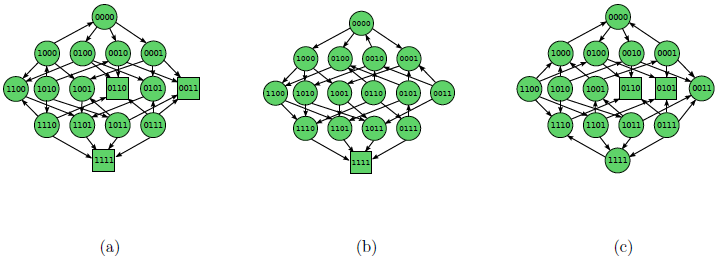
The evolutionary graphs for the fitness landscapes of *E. coli* with the antibiotics (a) Ampicillin (Amp), (b) Ampicillin + Sulbactam (Sam) and (c) Cefprozil (Cpr) for 4 possible substitutions found in *bla*_TEM-50_. Arrows represent fitness conferring mutations which can fix under the SSWM assumptions, the absence of an arrow in either direction corresponds to a neutral mutation which cannot fix under our assumptions. Squares denote local optima within the fitness landscapexs.

Mira et al. [2014] investigated the fitness landscapes of *E. coli* under 15 different β-lactam antibiotics using the mean minimum inhibitory concentration (MIC) of drug as a proxy for fitness for a total of N = 4 resistance conferring mutations. (Figure 2) Table Coding shows the evolutionary graphs of the fitness landscapes for three of these antibiotics, Ampicillin (Amp), Ampicillin+Sulbactam (Sam) and Cefprozil (Cpr). We will use these three fitness landscapes to demonstrate the steering hypothesis explicitly. In the case of a single peaked landscape, such as that for Ampicillin + Sulbactam, we cannot reduce the likelihood of resistance as all evolutionary trajectories lead to the global fitness optimum. It is only when a drug has a multi-peaked landscape that we may be able to avoid resistance through careful choice of preceding drugs. Of the 15 landscapes determined empirically by Mira et al. [2014] only the landscape for Ampicillin+Sulbactam is single peaked. In their review of empirical fitness landscapes de Visser and Krug [2014] find that biological landscapes show a variable but substantial level of ruggedness suggesting that multi-peaked landscapes could be quite common. Poelwijk et al. 2007, Poelwij 2011 showed that reciprocal sign epistasis within a landscape is necessary for the landscape to be multi-peaked.

**Fig. 3.**
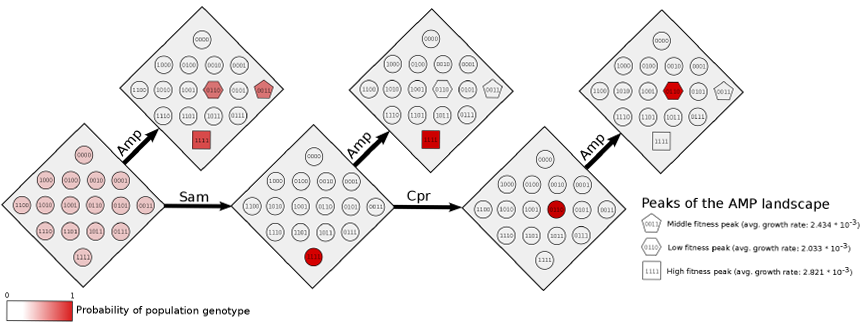
The probability distributions for accessibility of the peaks of the Amp landscape for different steering regimes. The starting distribution is μ = [1/2^N^,…, 1/2^N^]. When Amp is given first any of the three peaks of the landscape are accessible with most resistant genotype 1111 being most likely. If Sam is given first to steer the population to its sole peak 1111, then resistance to Amp will be guaranteed when it is applied. Alternatively, if Sam is given followed by Cpr then the population evolves to the local optima genotype 0110 of the Cpr landscape. If Amp is applied to this primed population the global optimum is inaccessible.

In the following we take *r* = 0 in (Equation 2) and note that changing the value of r will not change the accessibility of an evolutionary trajectory, hence by taking a different value of *r* ≥ 0 we will only change the result quantitatively (the probabilities may change) but not qualitatively. We begin by supposing that we do not know the initial population genotype. We can model this situation by taking as our prior population distribution μ = [1/2^N^,…, 1/2^N^] which specifies that each genotype is equally likely to constitute the starting population.

If we apply the drug Amp to this population distribution we find that in the expected distribution 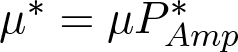 (shown in the first diamond in the top row of (Figure 3) Table Coding each of the fitness peaks could be found. In particular, the most highly resistant genotype 1111 can arise in the population with probability 0.62. Suppose instead we apply Sam first. In this case as the landscape is single peaked the population will converge to the global optimum genotype 1111. This genotype is also the global optimum of the Amp landscape and hence if we apply Amp after Sam we will encounter high resistance. We have steered the population with one drug to a configuration which increases the probability of resistance to a second. Next suppose instead we apply Cpr after Sam, in this case the population is guaranteed to evolve to a local optimum 0110 of the Cpr landscape. 0110 is the least fit local optimum of the Amp landscape. Thus if we apply Amp to the population primed by Sam and Cpr then evolution to the global optimum 1111 is not possible. This example demonstrates the steering hypothesis, that evolution can be shepherded through careful orderings of multiple drugs to increase or decrease the likelihood of resistance emerging.

To test our steering hypothesis we performed for each of the 15 drugs with landscapes derived by Mira et al. [2014] an *in silico* test of steering using combinations of one, two or three preceding drugs. Table 1 shows for each of the 15 antibiotics the combinations which reduce the probability of evolution to a peak fitness genotype to the lowest possible when applied in order followed by the final drug. We found that for 3 of the 15 drugs there exists another which steers an initial population μ to a configuration which prevents evolution to the global fitness optimum of the landscape entirely. This number rose to 6 when pairs of drugs applied sequentially are used to steer the population and to 7 when triples applied in sequence were considered.

We then performed a second *in silico* experiment to find combinations of steering antibiotics that maximize the probability that evolution proceeds to the least fit of the local optima of a final antibiotic. Table 3 show the results of this experiment. We found that, excluding the single peaked landscape for Ampicillin with Sulbactam, there exist 0 drugs for which a single other drug is able to steer the population to a configuration from which evolution to only the least fit optimum is possible. If pairs of drugs are used to steer there are 3 such drugs (including the example presented in the above demonstration) and if triples of steering drugs are considered there remains only 3. These findings suggest that through careful choice of steering drugs we may be able to prevent the emergence of resistance. During these experiments we found that 14/15 of the antibiotics in our experiment (Cefpodoxime (CPD) being excluded) appeared in an optimal steering combination of some length.

Whilst careful selection of drugs for steering can prevent the emergence of resistance, arbitrary drug orderings can also promote it. We performed an exhaustive *in silico* search of all singles, pairs of, and triples of steering drugs applied sequentially to prime the initial population for a final application of each of the 15 antibiotics. We found that steering with a single drug increased the likelihood of the most resistant genotype emerging in 57.3% of cases and decreased the likelihood in 29.8% of cases. Steering with pairs of drugs increased the likelihood in 64.1% of cases and decreased it in 28.4% of cases and steering with triples increased the likelihood in 65.6% of cases and decreased it in 27.5%. For each of the antibiotics except Cefaclor, Cefprozil and Ampicillin+Sulbactam (which is single peaked making steering irrelevant) we found that a random steering combination of length one, two or three is more likely to increase the chances of resistance than to reduce it. Indeed, for Piperacillin+Tazobactam and Ceftizoxime we found that a random steering combination will increase the probability of the most highly resistance genotype occurring in more than 80% of cases, suggesting that sequential multidrug treatments which use these antibiotics should proceed with caution. These findings suggest that the present system of determining sequential drug orderings without quantitative optimisation based guidelines could in fact be promoting drug resistance and that to avoid resistance we must carefully consider the order in which drugs are applied.

## Discussion

We have encoded Strong Selection Weak Mutation evolutionary dynamics on fitness landscapes as a Markov Chain. Through this encoding we can explore the dynamics of evolution by considering the algebraic properties of the associated Markov transition matrix. In particular, we have demonstrated that evolution on fitness landscapes is non-commutative through parallels with the non-commutativity of matrix multiplication and that antibiotic cycling strategies can be determined through matrix multiplication. We argue then that the ordering in which a collection of drugs is applied can significantly impact the population that exists after the application is complete.

We have shown that we can find sequences of drugs that can be applied to both avoid and promote the emergence of resistance in the population. In light of the slow pace of novel antibiotic discovery and the rapid emergence of resistance to the presently most utilized antibiotics, this finding suggests a new treatment strategy — one in which we use a sequence of drugs (or even treatment breaks which themselves impose a selective pressure [Andersson and Hughes, 2010]) to steer, in an evolutionary sense, the disease population to avoid resistance from developing. Further, the drugs used to prime the disease population for treatment by an effective antibiotic do not themselves need to be the most effective drugs available. This means that there could be a large pool of potential steering drugs in the form of antibiotics which have gone unused for many years due to their inefficacy.

**Table I.** For each of the 15 antibiotics we have derived the (ordered) sets of one, two and three steering drugs which reduce the probability of evolution to the maximal fitness genotype to the lowest possible. In the case that an ordered set of steering drugs reduced the probability to 0 we have not considered ordered sets of greater length (marked as – in the table). * — as the landscape for SAM is single peaked there can be no combination of steering drugs which reduce the probability of finding the global optimum. In all experiments the initial population distribution is taken as μ =[1/2^n^,…, 1/2^n^] and *r* = 0.

| Final drug | # of Peaks | Fittest peak genotype | Probability of ending at highest fitness genotype (no steering) | Best single steering drug | Probability of ending at highest fitness genotype (with best single steering drug) | Best ordered pair of steering drugs | Probability of ending at highest fitness genotype (with best steering pair) | Best ordered triple of steering drugs | Probability of ending at highest fitness genotype (with best steering triple) |
| --- | --- | --- | --- | --- | --- | --- | --- | --- | --- |
| Ampicillin (AMP) | 3 | 1111 | 0.62 | CPR | 0.25 | AMP→CPR | 0.0 | - | - |
| Amoxicillin (AMX) | 2 | 1101 | 0.75 | CTX | 0.62 | SAM→CPR | 0.51 | CRO→SAM→CPR | 0.51 |
| Cefaclor (CEC) | 3 | 0011 | 0.19 | SAM | 0.0 | - | - | - | - |
| Cefotaxime (CTX) | 4 | 1111 | 0.17 | CPR | 0.04 | AMP→CPR | 0.0 | - | - |
| Ceftizoxime (ZOX) | 2 | 0111 | 0.86 | AMX | 0.81 | SAM→AMX | 0.75 | CEC→CPR→AMC | 0.75 |
| Cefuroxime (CXM) | 2 | 0111 | 0.56 | AMC | 0.33 | CXM→AMC | 0.19 | AMC→CXM→AMC | 0.11 |
| Ceftriaxone (CRO) | 4 | 1111 | 0.28 | CEC | 0.20 | TZP→CEC | 0.05 | AMC→TZP→CEC | 0.0 |
| Amoxicillin +Clav (AMC) | 2 | 1101 | 0.67 | CXM | 0.39 | AMC→CXM | 0.23 | CXM→AMC→CXM | 0.13 |
| Ceftazidime (CAZ) | 3 | 0110 | 0.39 | TZP | 0.08 | AMC→TZP | 0.0 | - | - |
| Cefotetan (CTT) | 5 | 0111 | 0.18 | AMC | 0.0 | - | - | - | - |
| Ampicillin +Sulbactam (SAM) | 1 | 1111 | 1.0 | * | 1.0 | * | 1.0 | * | 1.0 |
| Cefprozil (CPR) | 3 | 0101 | 0.25 | AMP | 0.0 | - | - | - | - |
| Cefpodoxime (CPD) | 2 | 1111 | 0.81 | CRO | 0.74 | CTX→FEP | 0.57 | SAM→CEC→CRO | 0.50 |
| Piperacillin +Tazobactam (TZP) | 2 | 0101 | 0.80 | CTT | 0.72 | CTT→CTT | 0.72 | CTX→FEP→CTT | 0.68 |
| Cefepime (FEP) | 4 | 1111 | 0.48 | CTX | 0.39 | CAZ→CEC | 0.36 | CAZ→CEC→CTX | 0.33 |

**Table II.**
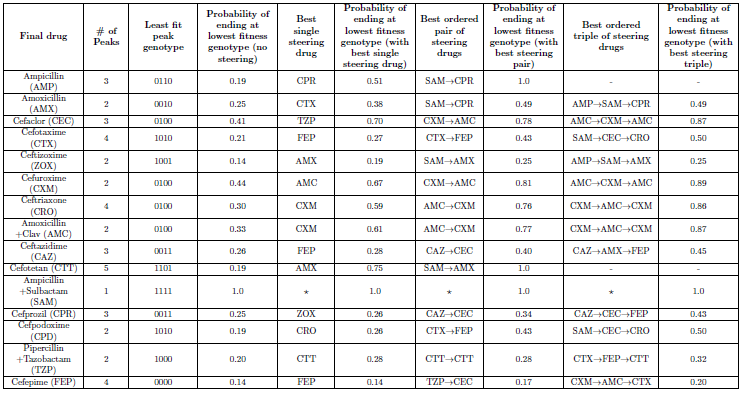
For each of the 15 antibiotics we determined how many of the possible single, ordered double and ordered triple combinations of steering drugs (allowing drugs to appear multiple times in the combination and also allowing the target drug to appear as a steering drug) improved or worsened the probability of the most resistant genotype being found after the steering drugs are applied in order followed by the final target drug. In each case the initial population was given by μ = [1/2^N^1/2^n^] and *r* = 0. As the SAM landscape is single peaked no combination of steering drugs will improve or worsen the outcome. As such, we have computed the overall numbers both with and without the contribution of the SAM row.

However, a major difficulty in using sequential drug treatments to steer disease populations is that in order to predict the outcomes we must know the fitness landscapes of the drugs involved. De Visser and Krug [2014] state that there exist less than 20 systematic studies of fitness landscapes and that these studies consider between 3 and 9 possible mutations. For steering to be fully effective we must account for all likely fitness conferring mutations and their effects on fitness under many drugs. Thus, many of the studies reviewed by de Visser and Krug are insufficient for determining clinically actionable steering strategies for certain diseases. Fortunately, for a number of highly resistant infectious diseases [Woodford and Ellington, 2007, Jensen and Lyon, 2009] and cancers [Lord and Ashworth, 2013, Lito et al., 2013] there are only a small number of mutations which seem to contribute to resistance. Further, recent work by Hinkley et al. [2011] in HIV has introduced a method to approximate large fitness landscapes from relatively fewer data points using a regression method. It follows that determining the landscapes is not an entirely intractable problem. A further complication in determining steering strategies is that fitness landscapes can be dependent on the disease microenvironment and have the potential to change from patient to patient or throughout the course of treatment. The consequences of such effects on fitness landscapes have not yet been experimentally determined.

Even in the absence of actual fitness landscapes our findings should be taken as a cautionary warning for multi-drug treatments, particularly those used to treat complex diseases such as multiple infections or cancers. In the same way that the drug ordering can be used to steer away from resistance we have shown it can also be used to make resistance more likely. Our results show that we may be inadvertently selecting for highly resistant disease populations through arbitrary drug ordering in the same way that highly resistant disease can emerge through irresponsible drug dosing. This result corroborates the findings of Pena-Miller and Beardmore [2010] that antibiotic cycling strategies can vary greatly in their efficacy and be both worse and better than mixing strategies. If we are to avoid resistance to our most effective drugs we must carefully consider how they are used together, both in combination and in sequence, with other drugs and take appropriate steps to reduce the risk.

Two major assumptions within our modeling are that drugs are administered for sufficiently long that evolution can converge to a local fitness optimum and that this convergence is guaranteed to occur. This assumption poses two potential problems in converting our model predictions to predictions of real-world bacterial evolution. The first is that if selection is strong and mutations are rare then there is a possibility of the population being driven to extinction before an adaptive mutation occurs. We have chosen to ignore this possibility within our modeling as in the context of treating bacterial infections this would constitute a success. The second is that the time to convergence could be prohibitively long for steering to constitute a realistic treatment strategy. We believe that the assumption of reasonable convergence times could be valid as adaptive walks in rugged landscapes are often short [Orr, 2005]. However, it has been shown that for certain landscapes there can exist adaptive walks of length exponential in the number of loci [Kaznatcheev, 2013], but since we get to choose those drugs with which to steer we can avoid landscapes for which the convergence time is prohibitively long. Further, our model is not necessarily restricted to the dynamics within a single patient. Goulart et al. [2013] used fitness landscapes to explore whole hospital scale antibiotic treatment strategies and our model, as an encoding of evolution on fitness landscapes, is capable of making predictions at this scale also. As such, even if evolutionary convergence is experimentally determined to be prohibitively slow for steering to be effective as a treatment for bacterial infection within a single patient, our results will still hold in scenarios which admit longer timescales. Specifically, evolutionary steering could provide an effective means to avoid the emergence of drug resistance at the hospital scale or in life-long diseases such as HIV.

The Strong Selection Weak Mutation model we have used here is a highly simplified, yet well studied model of evolution. The model ignores much of the complexity of the evolutionary process, specifically simplifying the genotype-phenotype map, ignoring the disease microenvironment and making the assumption of a monomorphic disease population in which deleterious and neutral mutations cannot fix. Under certain regimes of population size and mutation rates these simplifying assumptions break down. For example, if the population is sufficiently large then stochastic tunneling [Iwasa et al., 2004] - the situation where double mutations can occur allowing the crossing of fitness valleys - can arise causing a breakdown of the Strong Selection assumption. Similarly, if the mutation rate is sufficiently high then the population ceases to be monomorphic and forms a quasispecies [Nowak, 1992, Bull et al., 2005]. Conversely, if the population is sufficiently small then it becomes possible for deleterious mutations to fix [Moran et al., 1962, Wright, 1932, Fisher, 1958]. Finally, we have ignored the possibility of neutral spaces in the fitness landscape which have been shown to have significant impact on whether non-optimal genotypes can fix in the population as well as the time taken for evolution to find a locally optimal genotype [Schaper and Louis, 2014, Schaper et al., 2012]. We believe that each of these breakdowns of the SSWM model will have important consequences for the possibility and efficacy of steering and hence a proper treatment of their implications is beyond the scope of this paper. In our future work we aim to undertake a comprehensive study of the implications of population size, mutation rate, neutral drift and evolutionary convergence times on the steering hypothesis.

The SSWM model is a simplification of the evolutionary process and given that non-commutativity is present in this highly simplified model it is unlikely that commutativity will emerge as more complexity is introduced. It follows that the cautionary message regarding sequential drug application which results from our simplified model merits serious consideration. Whether or not measuring fitness landscapes provides sufficient information to correctly identify optimal drug orderings *in vivo* is a question that cannot be answered through mathematical modeling alone. It is only by verifying the predictions of steering strategies given by our model through biological experiment that we can determine whether they are viable. Supposing our model predictions are indeed viable then knowledge of some approximation to the fitness landscapes of the presently most used antibiotics could, in combination with our model, provide at least a good heuristic for how to proceed with multi-drug treatments, future antibiotic stewardship programs and clinical trial design.

**Table III.** For each of the 15 antibiotics we have derived the (ordered) sets of one, two and three steering drugs which increase the probability of evolution to the least fit optimum genotype to the highest possible. In the case that an ordered set of steering drugs increases the probability to 1 we have not considered ordered sets of greater length (marked as - in the table). * – as the landscape for SAM is single peaked there can be no combination of steering drugs which change the probability of finding the global optimum. In all experiments the initial population distribution is taken as μ = [1/2^N^,…, 1/2^N^] and *r* = 0

## Acknowledgment and Supplementary Information

We would like to offer the code underlying this model to any interested parties openly. DN would like to thank the Engineering and Physical Sciences Research Council (EPSRC) for generous funding for his doctoral studies. JGS would like to thank the NIH Loan Repayment Program for support. AGF is funded by the EPSRC and Microsoft Research, Cambridge through grant EP/I017909/1. ARAA and JGS gratefully acknowledge funding from the NCI Integrative Cancer Biology Program (ICBP) grant U54 CA113007 and they and PKM and RAG also thank the NCI Physical Sciences in Oncology Centers U54 CA143970 grant. We would also like to thank the Veterans Affairs Merit Review Program (RAB), the National Institutes of Health (Grant AI072219-05 and AI063517-07 to RAB), the Geriatric Research Education and Clinical Center VISN 10 (RAB), and an unrestricted research grant from the STERIS Corporation.

